# Encoding of odors by mammalian olfactory receptors

**DOI:** 10.1101/2021.12.27.474279

**Authors:** Aashutosh Vihani, Maira H. Nagai, Conan Juan, Claire A. de March, Xiaoyang S. Hu, John Pearson, Hiroaki Matsunami

**Affiliations:** Department of Neurobiology, Neurobiology graduate program, Duke University Medical Center, Durham, NC 27710, USA; Department of Molecular Genetics and Microbiology, Duke University Medical Center, Durham, NC 27710, USA; Duke Institute for Brain Sciences, Duke University, Durham, NC 27710, USA

## Abstract

1. Identified ligands for > 500 mouse ORs
2. ORs are specifically tuned towards individual odorants and their molecular properties
3. Odor molecular properties are informative of odor responses
4. Predictive modeling and convergent evolution analyses suggest specific residues within a canonical location for odorant binding

Olfactory receptors (ORs) constitute the largest multi-gene family in the mammalian genome, with hundreds to thousands of loci in humans and mice respectively^1^. The rapid expansion of this massive family of genes has been generated by numerous duplication and diversification events throughout evolutionary history. This size, similarity, and diversity has made it challenging to define the principles by which ORs encode olfactory stimuli. Here, we performed a broad surveying of OR responses, using an *in vivo* strategy, against a diverse panel of odorants. We then used the resulting interaction profiles to uncover relationships between OR responses, odorants, odor molecular properties, and OR sequences. Our data and analyses revealed that ORs generally exhibited sparse tuning towards odorants and their molecular properties. Odor molecular property similarity between pairs of odorants was informative of odor response similarity. Finally, ORs sharing response to an odorant possessed amino acids at poorly conserved sites that exhibited both, predictive power towards odorant selectivity and convergent evolution. The localization of these residues occurred primarily at the interface of the upper halves of the transmembrane domains, implying that canonical positions govern odor selectivity across ORs. Altogether, our results provide a basis for translating odorants into receptor neuron responses for the unraveling of mammalian odor coding.

## Introduction

Stimulus encoding and feature extraction are fundamental tasks performed by all sensory systems. Therefore, a central problem in neurobiology is defining how aspects of a stimulus are represented by the activity of sensory receptors^2–5^. This problem is particularly intriguing in the case of olfactory stimuli, which do not vary along a single, continuous dimension, such as wavelength or amplitude. Odorants, rather, have discrete molecular structures that determine their physical-chemical properties. An inability to relate how these discrete molecular structures and their associated physical-chemical properties influence receptor responses represents a major gap in knowledge. Consequently, one cannot robustly predict the neural activity patterns nor the perceptual attributes^6^ of an odorant starting from its physical-chemical properties.

A major hindrance in deciphering the coding of olfactory information by olfactory receptors (ORs) has been the historic inability to comprehensively identify ORs that respond to an odorant. Various *in vivo*, *ex vivo*, and *in vitro* methods have generally suffered from either a lack of insight into receptor identity or have been too low throughput for a comprehensive surveying of OR selectivity^5, 7–9^. With the mouse genome encoding over 1000 intact ORs, and odor reception following a combinatorial coding scheme, where one OR can be activated by a set of odorants and one odorant can activate a combination of ORs, defining a logic for peripheral odor coding is dependent on a comprehensive surveying while tracking receptor identity over a large odor panel^1, 10–12^.

Here, we performed a broad surveying of odorants *in vivo* to identify odorant-OR interactions in *Mus musculus*. By leveraging phosphorylated S6 ribosomal subunit capture (pS6-IP) coupled to RNA-Seq (pS6-IP-Seq), we were able to identify ORs expressed by recently active olfactory sensory neurons (OSNs; receptor deorphanization)^11, 13–15^. Then, using a library of molecular property descriptors, we parameterized the physical-chemical properties of the tested odorants to uncover relationships to the responses they elicited from cognate receptors. Finally, using our data, we asked 1) how well does odor molecular property similarity predict receptor response similarity and 2) if there are specific amino acid positions that influence odorant selectivity amongst receptors. Our results and analyses provide a foundational framework for understanding the molecular logic by which the quality of an odor molecule is encoded across a mammalian receptor repertoire.

## Results

### Estimation of chemical and receptor space sampling

First, we set out to identify ORs activated by a set of 61 odorants at various concentrations by leveraging pS6-IP-Seq. Immunoprecipitation of phosphorylated ribosomes from activated neurons followed by associated mRNA profiling by RNA-Seq, and differential expression analysis, enabled us to identify ORs expressed by OSNs activated by specific odorants (Supplementary figure 1A)^11, 14, 15^. ORs were considered odor-responsive if enrichment values (log_2_FC) were positive with a false discovery rate (FDR) < 0.05. Considering all odorants at all tested concentrations, this approach deorphanized a total of 555 ORs across 72 conditions (Supplementary table 1). Considering unique odorants yielding at least one activated OR at the lowest tested concentration, this approach deorphanized a total of 375 ORs across 52 odorants.

To examine the bias in our odorant set, we built an 1811-dimensional (1811D) space in which each dimension represented a molecular property descriptor^5^, such as molecular weight, number of atoms, or aromatic ratio, parameterizing the physical-chemical properties of the odor molecule. We then plotted our 52 uniquely tested odorants together with 4680 other small molecules^16^ other small molecules commonly found in foods and fragrances in this 1811D space to construct a chemical space consisting of a total of 4732 small molecules. Visualization of the first two principal components (PCs) did not reveal any obvious segregation of the test odorants, suggesting a broad sampling of chemical space by our test odor panel (Figure 1A, Supplementary figure 1B-C). To examine bias in our resulting deorphanized OR cohort, we computed pairwise OR Grantham distances^17^, an index of amino acid similarity, and visualized the results using multidimensional scaling (MDS). Examination of the first two MDS coordinates did not reveal any obvious segregation of the deorphanized 375 ORs, suggesting a broad sampling of receptor space (Figure 1B).

**Figure 1.**
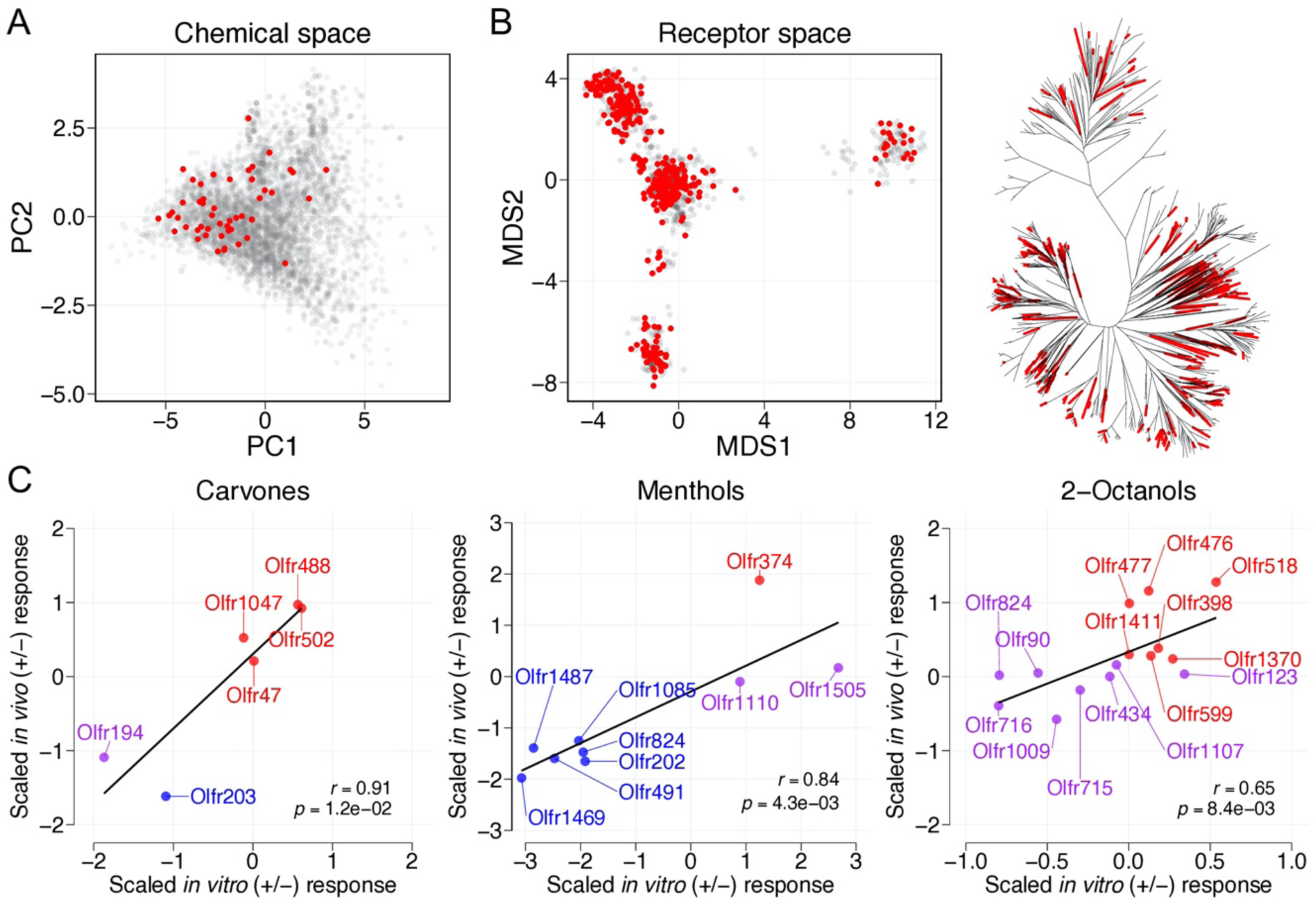
Data bias and pS6-IP-Seq validation. **A**, A total of 1811 molecular descriptors were calculated for a total of 4732 small molecules. The small molecules were projected onto a 2D chemical space made of the first and second principal components. The 52 unique odorants tested by pS6-IP-Seq at low concentrations, each yielding at least one activated OR, are colored in red. **B**, Left, Grantham distances were used to calculate a distance matrix for intact ORs. The matrix was visualized in two dimensions with multidimensional scaling to represent receptor space. ORs responding to at least one of the 52 tested odorants are colored red (*n* = 375). Right, a phylogenetic tree of intact ORs. Tree edges with identified and unidentified agonists at low concentrations are colored red and black respectively. **C**, ORs responsive to tested enantiomers were evaluated by heterologous expression. ORs enriched by pS6-IP-Seq against (+)-odorant are colored red while ORs enriched by (-)-odorant are colored blue. ORs enriched by both enantiomers are colored purple. Linear regression reveals *in vivo* and *in vitro* responses to be highly correlated (carvones *r* = 0.91, *p* = 0.012; menthols *r* = 0.84, *p* = 0.0043; 2-octanols *r* = 0.65, *p* = 0.0084).

To validate the receptor specificity of the pS6-IP-Seq dataset, we selected enantiomers (carvones, menthols, and 2-octanols) for *in vitro* testing. We transiently expressed ORs responsive to tested enantiomers in Hana3A cells and challenged with individual odorants to generate dose response curves. Comparison of *in vitro* responses to *in vivo* responses revealed the data to be highly correlated (carvones *r* = 0.91, *p* = 1.2E-2; menthols *r* = 0.84, *p* = 4.3E-3; 2-octanols *r* = 0.65, *p* = 8.3E-3). Altogether, these results substantiated the pS6-IP-Seq dataset and yielded confidence that the pS6-IP-Seq strategy would provide an index of receptor selectivity even amongst structurally similar odorants (Figure 1C, Supplementary figure 2A-F).

### Describing receptor tuning

Having broadly sampled chemical and receptor spaces, we next sought to quantify the relative responses of individual receptors to the test odor panel. Individual receptors displayed unique response profiles across the odorants. Examining receptor tuning did not reveal a bimodal distribution of narrowly and broadly tuned receptors, but rather a continuum of tuning breadths with an average of 1.85 cognate odorants per significantly responding receptor (Figure 2A-D, Supplementary figure 3A-B).

**Figure 2.**
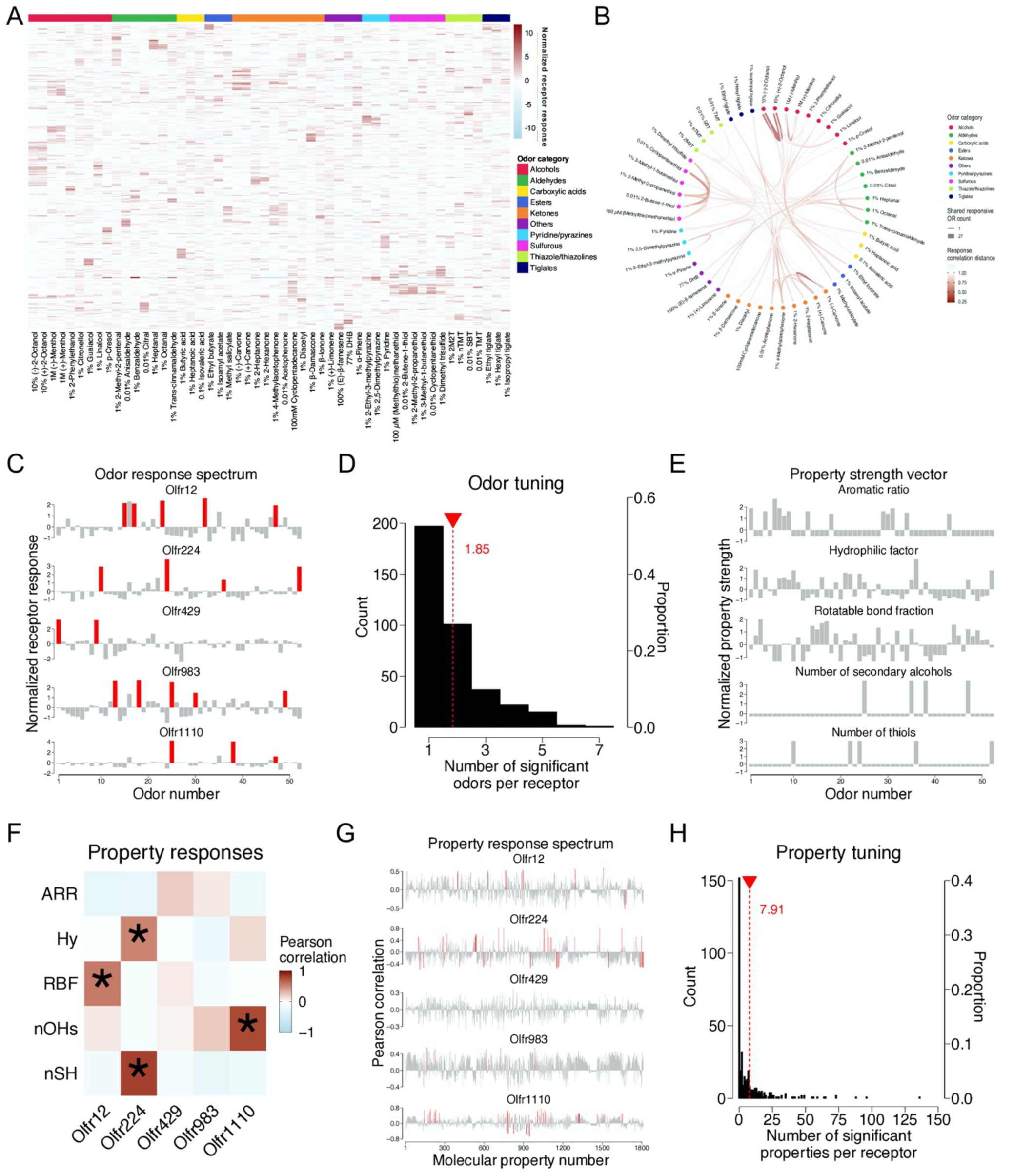
Combinatorial coding and OR tuning. **A**, Odor responses across the 52 unique odorants, tested at low concentrations, visualized by heatmap. A total of 375 ORs were determined to be responsive to at least one of the tested odorants (log_2_FC > 0 and FDR < 0.05). Responses are normalized such that each odor has zero response mean and one standard deviation (z-scored). Responses are color coded from negative, to zero, to positive responses in light blue to white to red. The following odorants are abbreviated: 2-methyl-2-thaizoline (2M2T), 2,4,5-trimethyl-4,5-dihydrothiazole (TMT), 2-*sec*-butyl-4,5-dihydrothiazole (SBT), 2,4,5-trimethylthiazole (nTMT), and 3,4-dehydro-exo-brevicomin (DHB). ORs were sorted by correlation distance. Odorants were sorted by odor category. **B**, Odor responses across the same 52 odorants visualized by chord plot. Odorants were sorted by odor category and associations were visualized. Association band thickness corresponds to the number of ORs shared, while color corresponds to overall response similarity across the 375 deorphanized ORs using correlation distance (1-*r*). **C**, Z-scored odor response spectra of five example ORs across the 52 tested odorants. Responses were normalized such that each OR is z-scored. **D**, Histogram of the number of significant odorants per receptor. On average, each responding OR was activated by 1.85 odorants (*n* = 692). **E**, Z-scored PSVs of five example molecular properties across the same set of odorants as B. **F**, Property responses given by the Pearson’s correlation coefficients between the odor responses (B) and PSVs (D) calculated over the 52 odorants in the panel. The following molecular properties are abbreviated: aromatic ratio (ARR), hydrophilic factor (Hy), rotatable bond fraction (RBF), number of secondary alcohols (nOHs), and number of thiols (nSH). **G**, Property response spectra characterized by Pearson’s correlation coefficients between the five example ORs and 1811 molecular properties. **H**, Histogram of the number of significant molecular properties per receptor (*n* = 2967). On average, each odor-responsive OR was significantly correlated with 7.91 molecular properties with a median of 2 (FDR < 0.05). Out of the 375 deorphanized ORs, 223 ORs displayed significant correlations to at least one molecular property descriptor.

To describe the tuning of ORs towards specific molecular properties, we next generated property strength vectors (PSVs) for each of the molecular descriptors (Figure 2E)^18^. The responsiveness of each OR to each molecular property was then characterized as a Pearson’s correlation between the odor response spectrum and the values taken by the PSV across the 52 odorants tested (Figure 2F). The array of such correlations (hereby termed property response spectrum) taken across the molecular property descriptor set defined the molecular receptive range and property tuning of each OR (Figure 2G). For example, several ORs that displayed robust responses towards thiol odorants yielded tuning towards the “number of thiol groups” molecular property (Supplementary figure 4A-D). The number of properties that single receptors responded to significantly (FDR < 0.05) varied from receptor to receptor with a range of 0 to 136 (mean = 7.91, median = 2). Indeed, the majority of deorphanized ORs (223/375) displayed significant correlations to at least one of the molecular property descriptors (Figure 2H). Within the subset of significant OR response-property pairs (2967/ 679125), correlations spanned both negative (-0.77 to -0.49) and positive (0.49 to 0.82) values with an absolute average of 0.55 (Supplementary figure 3C).

### Odor molecular properties are informative of odor response patterns

Having identified receptor responses to a large and diverse set of odorants, we next sought to determine the effectiveness of using odor molecular properties to predict receptor responses via similarities^5, 18, 19^. To describe the similarity between odorants in molecular property space, we calculated distances of normalized property strength values between odorant pairs. To represent odor similarity in OR response space, we similarly calculated pairwise distances between normalized receptor responses. Linear regression between odorant similarity distances and response similarity distances revealed a significant relationship (*r* = 0.29, *p* = 3.4E-27; Figure 3A, Supplementary figure 5A-B), implying odor molecular properties were informative of receptor response patterns.

**Figure 3.**
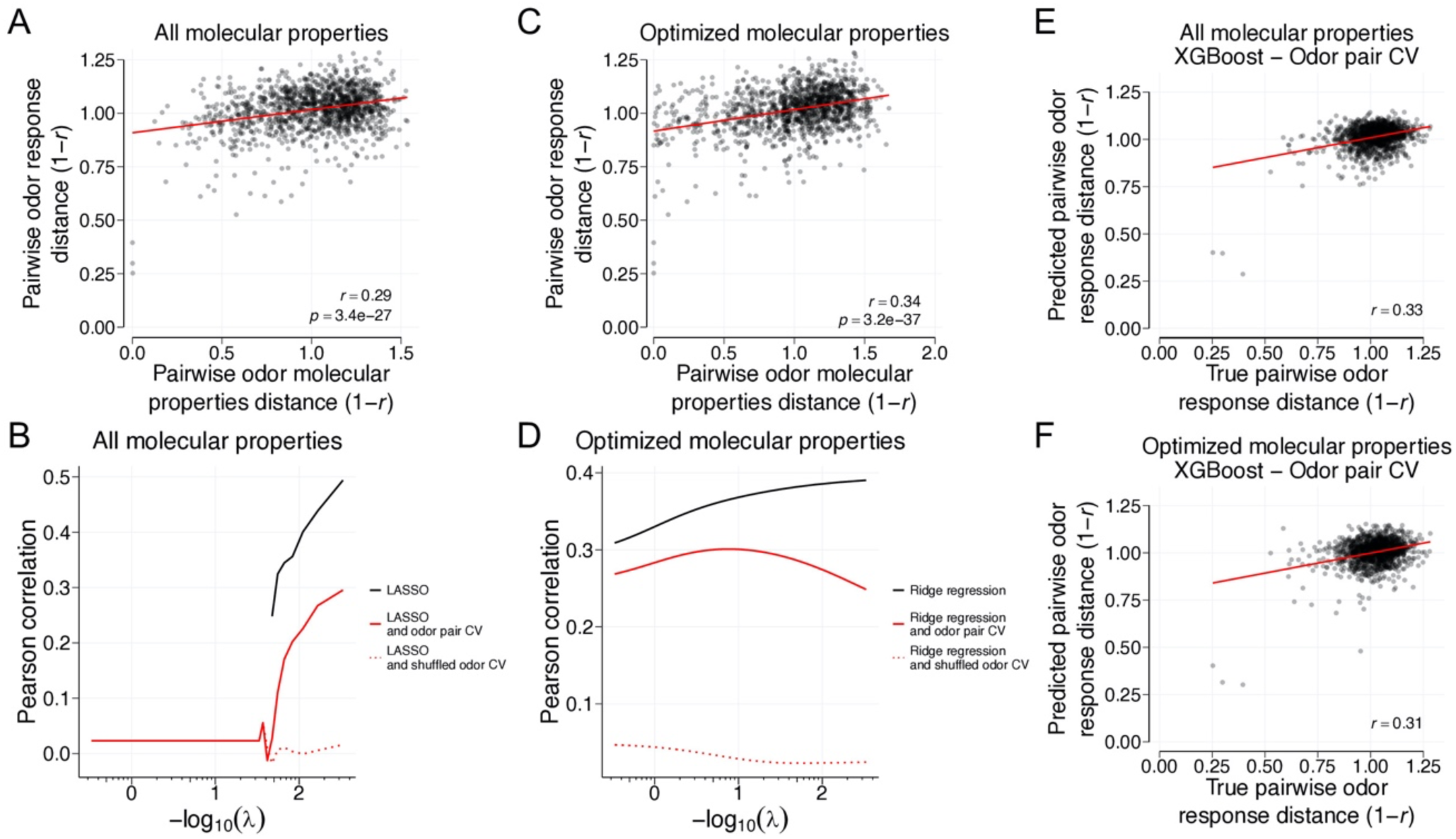
Pairwise odor similarity comparisons in response and property spaces. **A**, Pairwise correlation distance measurements between odorants in response space regressed against molecular property space (*r* = 0.29, *p* = 3.4E-27, *n* = 1326). **B**, LASSO regression as a function of varying the loss function (λ). Large λ values (leftmost edge) generally correspond to the selection of a few weighted molecular properties. Small λ values (rightmost edge) generally correspond to the selection of many weighted molecular properties. By increasing the number of weighted molecular properties selected by the LASSO algorithm, odor responses become increasingly well fit by molecular properties (black line). To evaluate the generalizability of the sparse property selection, pairs of odorants were iteratively held-out during training and added back intact (solid red line) or shuffled (dashed red line) for cross-validation. Euclidean distances were used to quantify differences between pairs of odorants in molecular property space. Correlation distances were used to quantify differences between pairs of odorants in response space. **C**, Pairwise correlation distance measurements between odorants in response space against optimized molecular property space (*r* = 0.34, *p* = 3.2E-37, *n* = 1326). **D**, Ridge regression results as a function of varying the λ loss function using the optimized set of molecular properties. Large λ values (leftmost edge) generally correspond to dependence on a few weighted molecular properties. Small λ values (rightmost edge) generally correspond to dependence on many weighted molecular properties. **E**, Results of odor pair cross-validation using the XGBoost model framework with default hyperparameters with all molecular properties (*r* = 0.33, *n* = 1326). **F**, Results of odor pair cross-validation using the XGBoost model framework with default hyperparameters with the optimized set of molecular properties (*r* = 0.31, *n* = 1326).

We next considered the possibility that a subset of the molecular property descriptors may be better able to relate odor molecular property similarities to receptor response similarities. To test this possibility, we built a sparse regression and performed feature selection using the Least Absolute Shrinkage and Selection Operator (LASSO). By varying the LASSO loss function (λ) to influence the number and relative contribution of the selected molecular properties, we observed improved correlations with increasing numbers of weighted molecular properties. Importantly, by performing odor pair cross-validation, we observed that parsimonious combinations of odor molecular properties selected by LASSO yielded positive predictive abilities (optimal correlation distance odor pair cross-validation *r* = 0.30, Figure 3B, Supplementary figure 6A-B). To complement these findings, we also selected an “optimized” set of 65 molecular properties; which included descriptions of aromaticity, functional group, and molecular geometry; that could be individually linearly decoded and regressed from OR response patterns alone (Supplementary figure 7A, Supplementary table 2). Using ridge regression with the “optimized” set of 65 molecular properties, we could again predict response similarities from molecular property similarities (optimal correlation distance odor pair cross-validation *r* = 0.30, Figure 3C-D, Supplementary figure 7B-C). Altogether, we interpreted these “optimized” set of 65 molecular property descriptors as both, being capable of explaining OR response variance, and contributing to the natural statistics of odorants.

To further validate the predictive abilities of molecular properties and our molecular property optimization, we also trained and cross-validated a feed-forward non-linear model (XGBoost). In the first cross-validation scheme, we performed odor-pair cross-validation using all calculated molecular properties as predictors. In the second, we limited molecular properties to the “optimized” set. In both cross-validation schemes, predicting response similarities from molecular properties outperformed shuffled controls (Figure 3E-F, Supplementary figure 8A-B). Altogether, these results show that odor responses can be explained in part by combinations of molecular property descriptors.

### Specific receptor residues predict ligand selectivity of ORs

Comprehensive identification of ORs responsive to many odorants prompted us to next search for generalizable relationships between odorants, ORs, and receptor residues^11, 14^. To do so, we built logistic models using aligned ORs. For each odor fit with a regularized logistic model, receptors were randomly split into 90% training and 10% testing sets for 100 repetitions. Iterating this process over the set of tested odorants identified a series of weighted positions, harboring amino acids with predictive power, occurring primarily at the upper halves of the fourth and fifth transmembrane domains (TMDs) (Figure 4A). For example, visualizing amino acids occurring at these positions identified an enrichment of cysteine and methionine residues in TMD5 amongst ORs responding to sulfurous odorants (Supplementary figure 9A). Regressing the average weight assigned to each position, from odorants solvable by logistic regression (area under receiver operating characteristic curve, AUROC > 0.5), by percent conservation revealed an anti-correlation (*r* = -0.38, *p* = 6.1E-12, Figure 4B-C)^1^. Using a Support Vector Machine (SVM) classifier, with a linear kernel, led to similar predictions regarding the response likelihoods of held-out ORs (Logistic regression AUROC = 0.70, Linear SVM AUROC = 0.78, Figure 4D, Supplementary figure 10A).

**Figure 4.**
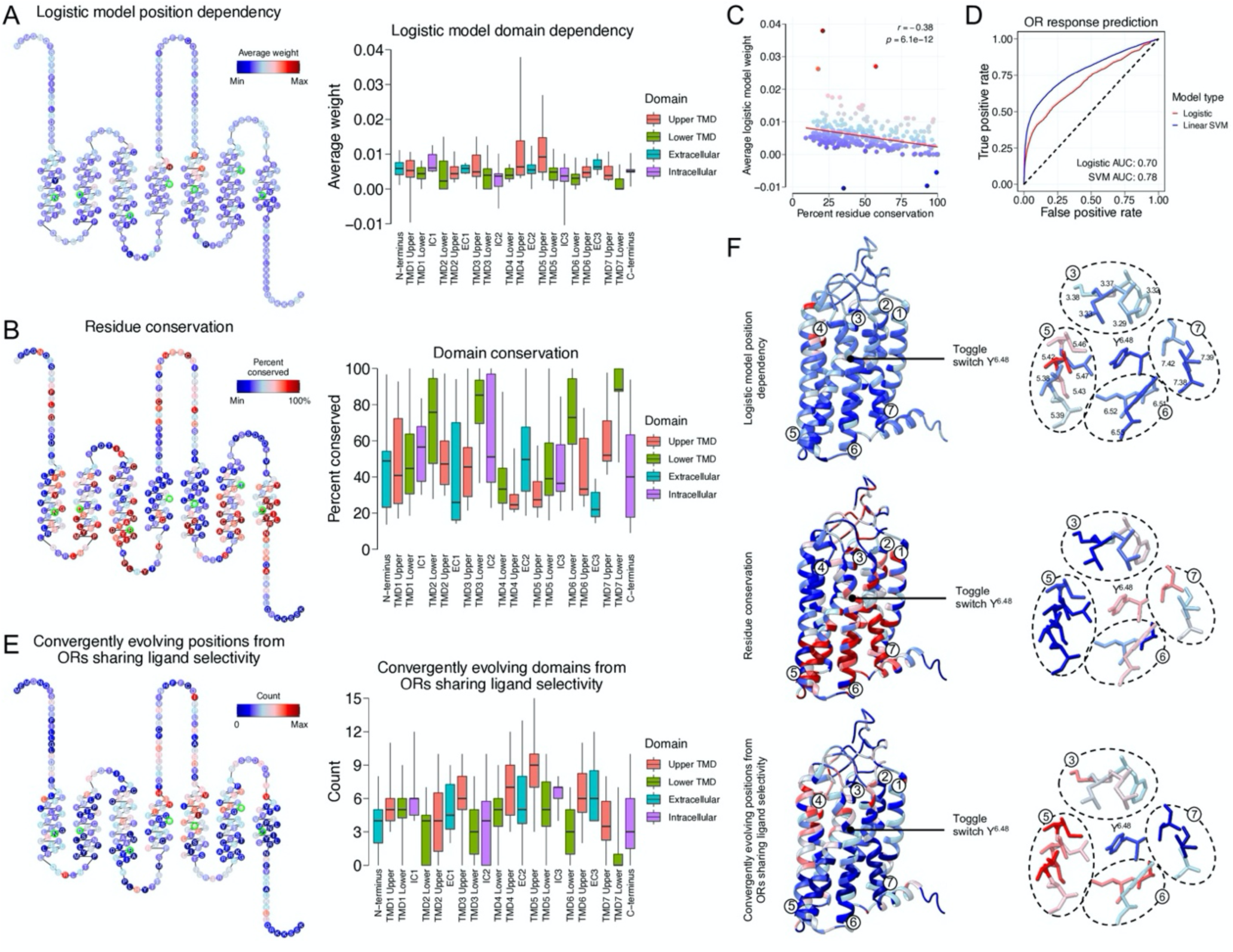
Sequence-function relationships of ORs. **A**, ORs were split into 90% training and 10% testing sets for 100 repetitions of regularized logistic regression using aligned sequences as predictors. Odorants which yielded AUROC values > 0.5 (Supplementary table 3) were subset and non-zero weights were averaged across positions, repetitions, and odorants. Individual weights were visualized by snakeplot (left) with the most commonly occurring amino acid described at each position. Ballesteros-Weinstein x.50 numbers for each TMD are highlighted in green. Weight distributions were visualized across domains by box-and-whisker plot (right). **B**, Left, the most commonly occurring amino acid was quantified by its percent conservation across ORs and visualized by snake plot. Right, percent conservation distributions were visualized across receptor domains by box-and-whisker plot. **C**, Regressing the average weight assigned to a position, by regularized logistic regression, by its percent conservation reveals an anti-correlation (*r* = -0.38, *p* = 6.1E-12). **D**, Using a SVM classifier with a linear kernel to predict response likelihoods of held-out ORs leads to similar results as logistic regression (logistic regression AUROC = 0.70, linear SVM AUROC = 0.78). **E**, ORs sharing response to an odorant were subset and pairwise compared to a null set of ORs with comparable protein sequences. Pairwise Grantham distance distributions between the responsive and null sets were compared by Kolmogorov-Smirnov statistical test to determine if amino acid similarity distributions were different. Count measurements reflect the number of times a position harbored amino acids with differing distributions between the two groups at an FDR < 0.05. Left, these results were visualized by snakeplot. Right, these results were visualized by box-and-whisker plot describing domain distributions. **F,** Left, visualization of the data in 3D using homology models. Color schemes are identical those in panels A, B, and E. TMDs are indicated. Right, zoomed in view focusing on residues found directly above the conserved Y^6.48^ toggle switch residue. A consensus of poorly conserved residues found in TMD3, 5, and 6 can be seen to exhibit both higher weights by regularized logistic regression models, and convergent evolution. Ballesteros-Weinstein numbers associated with displayed residues are also described.

The massive expansion and rapid evolution of the OR gene family posits opportunities for the convergent evolution of distantly related ORs to evolve odorant selectivity independently. To search for receptor sequence positions exhibiting convergent evolution, we asked if ORs sharing response to an odorant possessed positions harboring amino acids with physical-chemical properties, measured by Grantham’s distance^17^, which deviated from comparable but odor-unresponsive ORs. Iterating over the set of tested odorants, this analysis identified a series of poorly conserved positions (*r* = -0.54, *p* = 6.5E-25, Figure 4E, Supplementary figure 10B), especially localized to the upper half of TMD5. Regressing the average weight assigned to each position, via regularized logistic regression, by the number of times each position displayed convergent evolution, revealed similar findings between the two approaches (*r* = 0.42, *p* = 1.5E-14, Supplementary figure 10C). Importantly, the localization of these positions, and those identified by logistic models, was consistent with a region implicated in ligand binding in other class A GPCRs^20–25^. Altogether these results show the odorant selectivity of ORs are in part explained by convergently evolving residues occurring at a common site of poorly conserved residues within the TMDs.

To visualize the results of our analyses in 3D, we next built an OR homology model. Focusing on the conserved “toggle switch” Y^6.48^ residue previously reported to reside at the bottom of the ligand-binding cavity of other class A GPCRs^20, 26–28^, we consistently observed nearby residues in the upper halves of TMD3, TMD5, and TMD6 as exhibiting heavy weights in our logistic models, poor conservation, and convergent evolution, implying a canonical cavity for odorant binding across our tested odorants. Altogether, these results are consistent with the idea that few mutations within the ligand binding site of ORs can broadly reconfigure chemical tuning, a feature that is likely to have facilitated the rapid evolution of receptors with distinct ligand specificities.

## Discussion

Given the inordinate complexity of the chemical world, large repertoires of ORs appear to be necessary for the detection and discrimination of diverse chemicals in the environment, as exemplified by the significant genomic space that is systematically subjugated to ORs across numerous species. These findings are compounded by the identification of OR-specific chaperone proteins which may allow functional expression of ORs with cryptic mutations, further underscoring the high degree of sequence diversification ORs are enabled to possess^29, 30^. Using a diverse set of odorants, here we have performed a functional *in vivo* characterization of the OR repertoire of *Mus musculus*. Linking the activity of the receptor repertoire to an extensive set of molecular property descriptors parameterizing the physical-chemical properties of the odorants, we learned that ORs displayed a continuum of tuning breadths. Similarities between sparse sets of molecular properties could be used to predict receptor response patterns. Finally, analyses linking odorant selectivity and amino acid residues most consistently identified a series of poorly conserved residues located primarily in the upper half of the transmembrane domains.

While our test odor panel, and resulting deorphanized receptor set, was broad in the coverage of odor and receptor spaces, the data was by no means all-encompassing. 20 PCs were required to cover 90% of variance in the 52 set of tested odorants, whereas the full set of 4732 small molecules required 74 PCs to achieve comparable coverage (Figure 1 – Supplementary figure 1B). Our analyses therefore likely reflect lower bound estimates of chemical and receptor response spaces. Nevertheless, these results also imply that a substantial amount of the information in the molecular property descriptors is highly redundant. Furthermore, we note that the dimensionality of the tested odorants in receptor response space is higher than the dimensionality of the odorants in chemical space, with 34 PCs needed to explain more than 90% of the response variance (Figure 1 – Supplementary figure 1B). This increased dimensionality indicates there are facets of odor response by ORs that are poorly explained by a similar number of flat surfaces in chemical space described by molecular property descriptors. Although the molecular property descriptors used in this study can explain and predict response similarities, further searches for latent descriptors capable of better associating odor properties to their receptor responses may improve these associations^31, 32^.

Several implications arise from our observation that diverse odorants share commonalities that relate their receptor responses to amino acid residues. First, the poor conservation of positions harboring residues exhibiting predictive power and convergent evolution suggest a mechanism by which flexible chemical recognition can be achieved by a family of proteins while maintaining a degree of conservation necessary for functional protein integrity and activation of conserved downstream signaling cascades. Second, the association of the third, fifth, and sixth transmembrane domains with odor selectivity are also consistent with site-directed mutagenesis efforts on single ORs that have been shown to influence OR responses^33–37^. Finally, these results are consistent with recent evidence from structural elucidation of an ionotropic insect OR, which revealed a single binding pocket for a structurally diverse odorants^38^. Future structural elucidation of mammalian ORs will enable direct addressing of the modes of odorant-OR interactions.

In summary, we have provided a systematic, quantitative analysis of the primary representation of an odor, as registered by the differential responses of individual ORs. Our results and analyses provide a foundational framework for investigating how these primary odorant representations are transformed into subsequent representations to ultimately guide behavioral outputs.

## Supplementary figures

**Supplementary figure 1.**
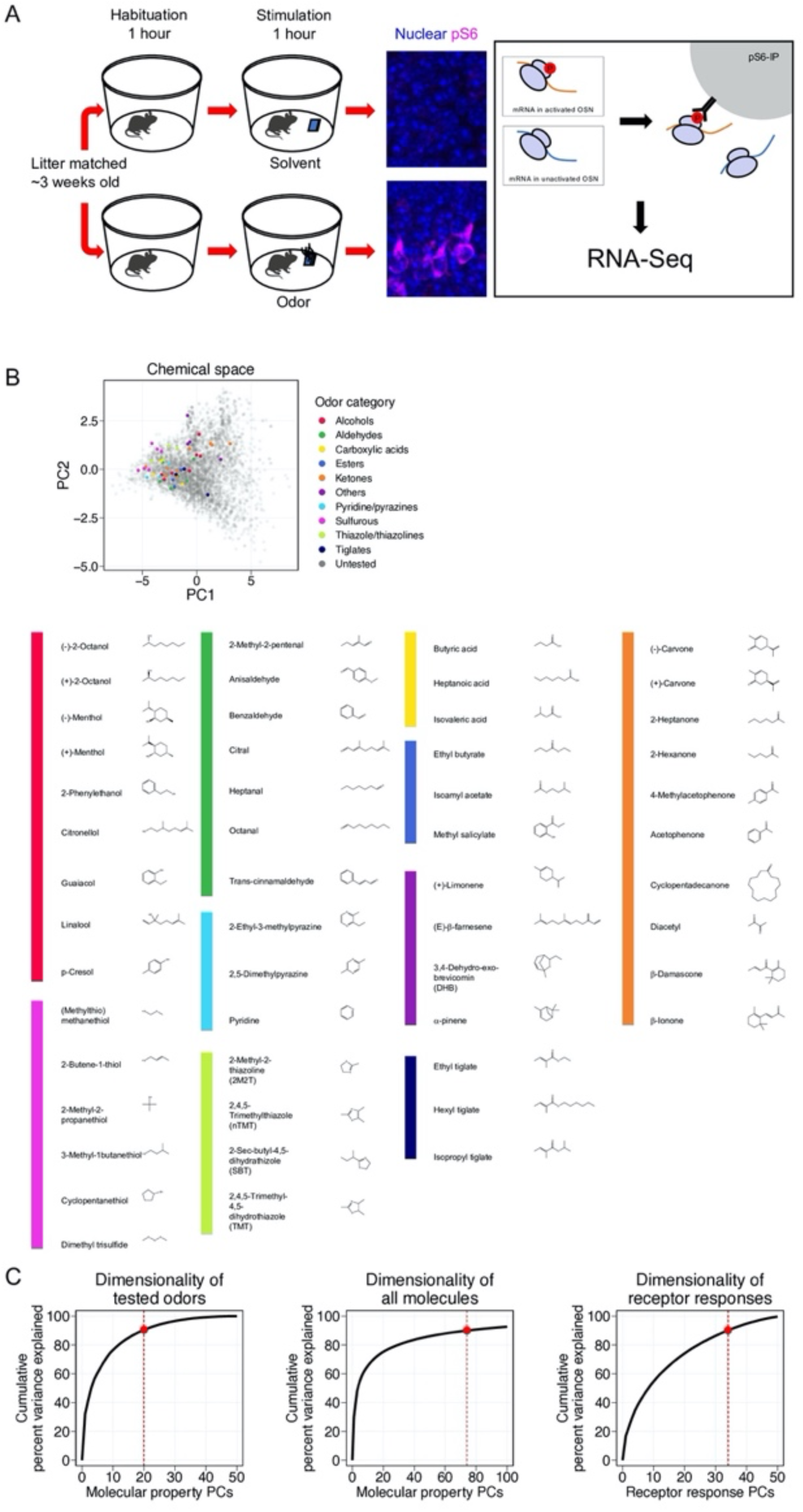
Details of chemical and receptor space sampling. **A**, Schematic of the pS6-IP-Seq experiment. Litter matched, ∼3 weeks old mice were used. Mice were first habituated to an odor-free environment for 1 hour. One mouse then received exposure to an odor stimulus, while another received exposure to the solvent, again for 1 hour. Whole olfactory mucosa was then harvested and immunoprecipitated using an antibody against pS6 and subjected to RNA-Seq. **B,** Top, tested odorants in chemical space colored by odor category. Bottom, molecular structures of tested odorants sorted by odor category. **C**, Left, the cumulative percent variance of the tested odorants in chemical space explained as a function of included PCs of molecular properties. A minimum of twenty PCs was required to capture at least 90% of the variance for the test odorants in chemical space. Middle, the cumulative percent variance of all molecules in chemical space explained as a function of included PCs of molecular properties. A minimum of seventy-four PCs was required to capture at least 90% of the variance for all molecules in chemical space. Right, the cumulative percent variance of the tested odorants in response space explained as a function of included PCs of receptor responses. A minimum of thirty-four PCs was required to capture at least 90% of the variance for the test odorants in response space.

**Supplementary figure 2.**
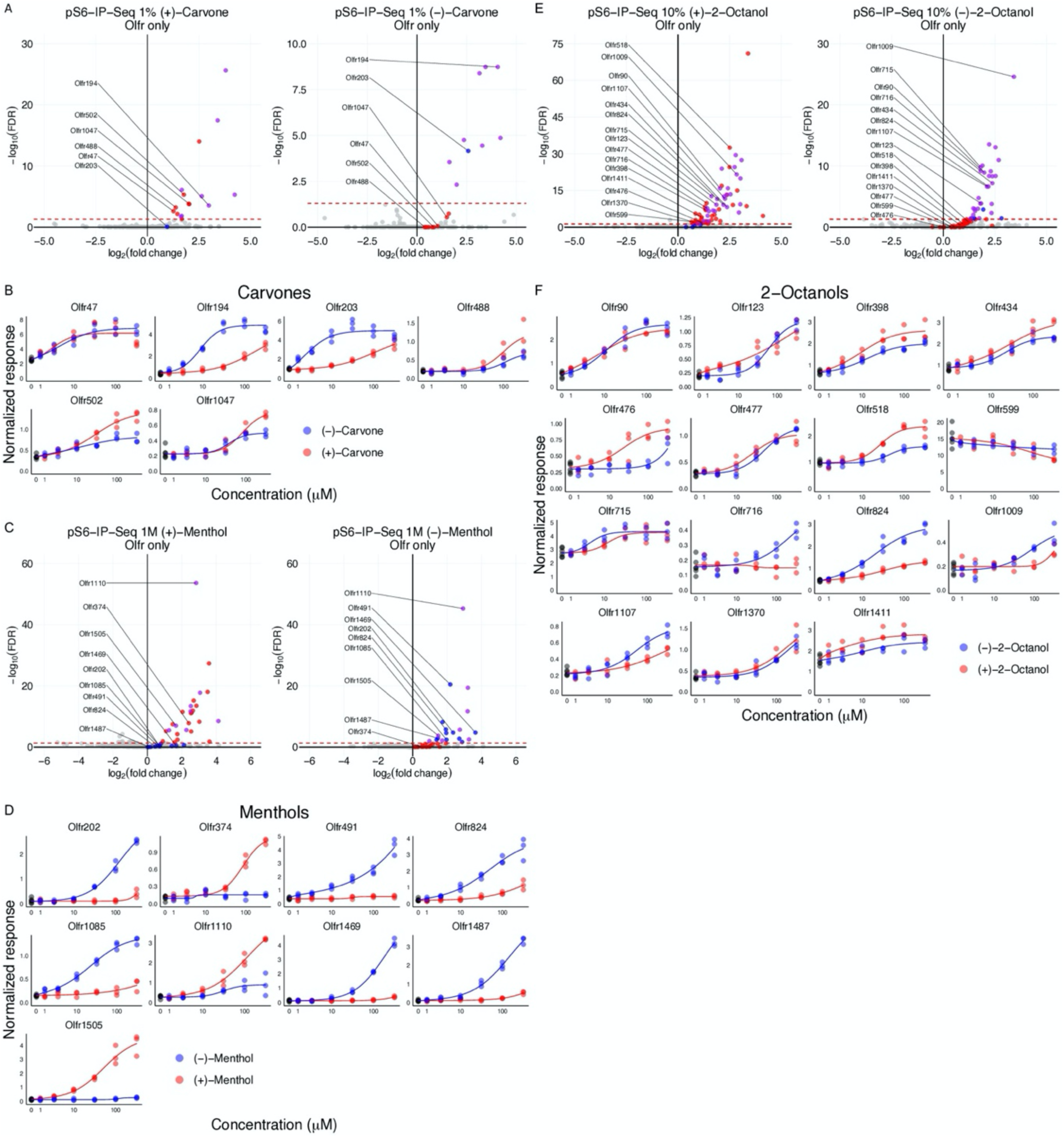
Details of *in vitro* validation. **A**, Volcano plots for pS6-IP-Seq results by exposure of mice to 1% (+)-carvone and 1% (-)-carvone. A red line at FDR = 0.05 is drawn. ORs enriched by just (+)-odorant are colored red while ORs enriched by just (-)-odorant are colored blue. ORs enriched by both enantiomers are colored purple. Labeled ORs were validated *in vitro*. **B**, Dose response curves of ORs displaying *in vitro* response to at least one of the tested carvone enantiomers. **C**, Volcano plots for pS6-IP-Seq results using 1M (+)-menthol and 1M (-)-menthol. **D**, Dose response curves of ORs displaying *in vitro* responses towards menthol enantiomers. **E**, Volcano plots for pS6-IP-Seq results using 10% (+)-2-octanol and 10% (-)-2-octanol. **F**, Dose response curves of ORs displaying *in vitro* responses towards 2-octanol enantiomers.

**Supplementary figure 3.**
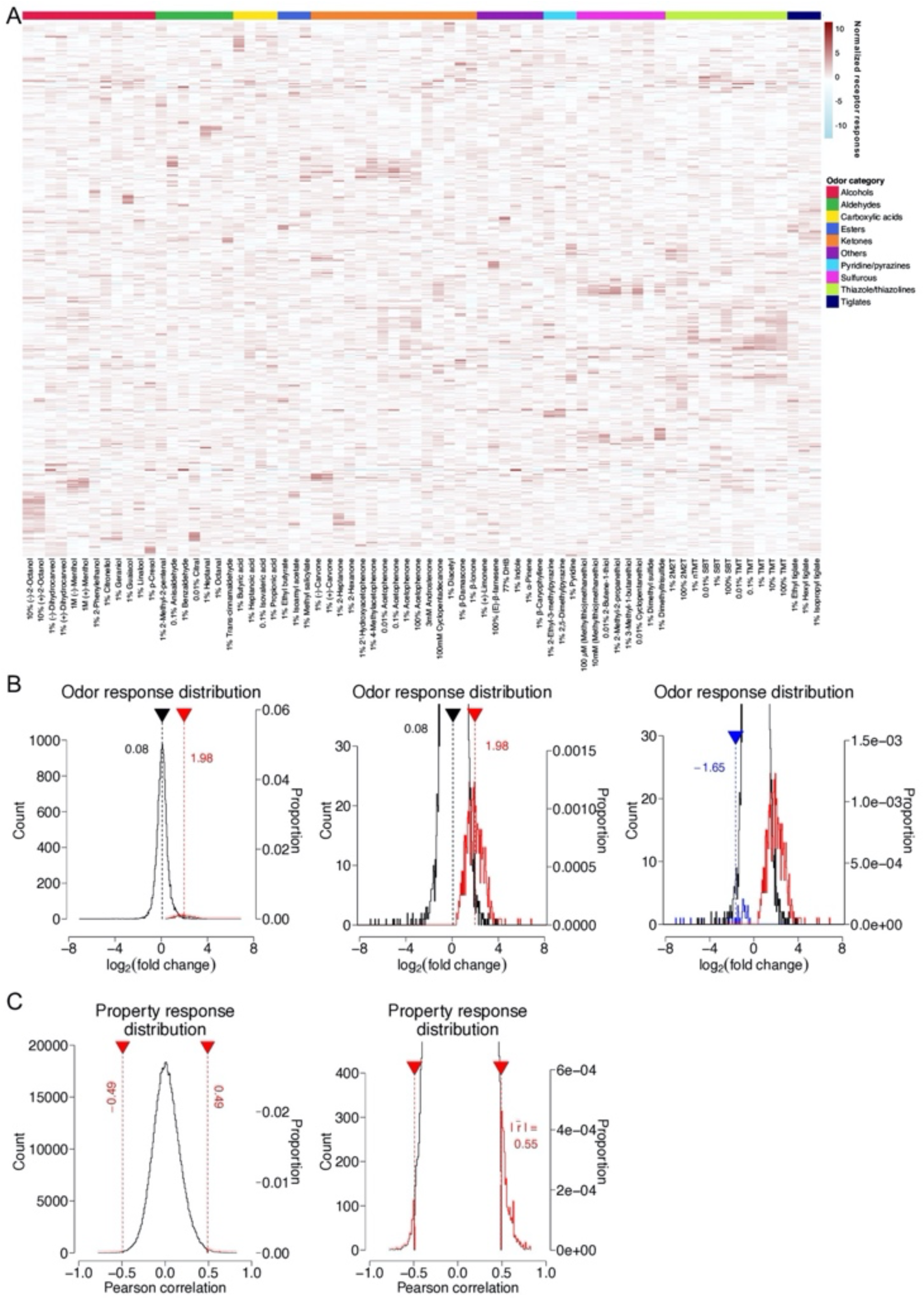
Details of OR response properties. **A**, Odorant responses determined by pS6-IP-Seq across all tested 72 odorants/concentrations visualized by heatmap. A total of 555 ORs are determined to respond to at least one of these odorants/concentrations. Responses are z-scored by odor. The following odorants are abbreviated: 2-methyl-2-thaizoline (2M2T), 2,4,5-trimethyl-4,5-dihydrothiazole (TMT), 2-*sec*-butyl-4,5-dihydrothiazole (SBT), 2,4,5-trimethylthiazole (nTMT), and 3,4-dehydro-exo-brevicomin (DHB). Odorants are sorted by functional group while ORs are sorted using correlation distance. **B**, Left, histogram (bin size = 0.05) of the distributions of OR responses across the panel of 52 odorants tested at low concentrations. Significant OR-odor pairs (log_2_FC > 0 and FDR < 0.05) are colored red, while non-significant pairs are colored black. OR-odor pairs that were classified as nonsignificant had an average log_2_FC enrichment of 0.08 by pS6-IP-Seq. OR-odor pairs classified as significant had an average log_2_FC enrichment of 1.98 by pS6-IP-Seq (nonsignificant responses *n* = 18808, significant responses *n* = 692). Middle, zoomed in. Right, A small number of inhibitory responses (log_2_FC < 0 and FDR < 0.05) were observed in the data. These responses were otherwise classified as nonsignificant (*n* = 44). **C**, Left, distribution of the OR-molecular property pairwise Pearson correlation coefficients using the 52 odorants tested at low concentrations (bin size = 0.01). At an FDR < 0.05, the Pearson correlation coefficient cutoffs were -0.49 and 0.49 for negative and positive correlations respectively, with an absolute average of 0.55 for significant correlations (nonsignificant correlations *n* = 676158, significant correlations *n* = 2967). Right, zoomed in.

**Supplementary figure 4.**
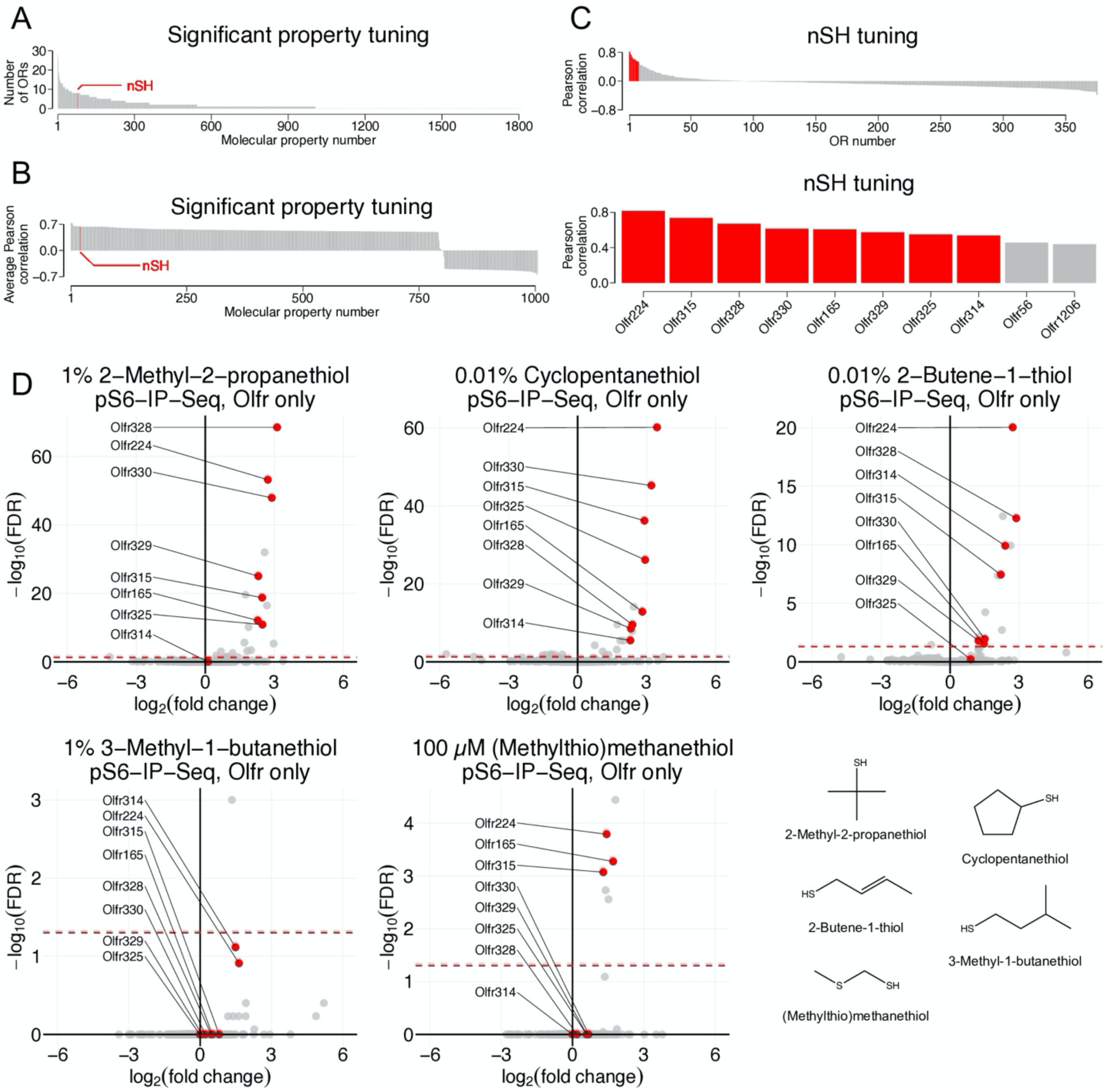
Details of thiol property tuning. **A,** Tuning towards molecular properties described by the number of significant OR-molecular property associations. Out of the 1811 molecular properties, 1005 were tuned towards by at least one OR. Molecular property nSH (number of thiol groups) is highlighted in red. **B**, Visualization of the average Pearson correlation tuning value for ORs that were significantly tuned toward each of the 1005 molecular properties. **C**, Top, tuning of individual ORs, measured by Pearson correlation, towards nSH. Bottom, the top ten ORs displaying nSH tuning. Eight of these ten ORs were significant (FDR < 0.05). **D**, Raw pS6-IP-Seq differential expression data, visualized by volcano plot, with chemical structures of the tested thiol odorants. ORs that were tuned towards thiol are highlighted in red. A consensus of thiol odorant response can be observed for the ORs tuned towards nSH.

**Supplementary figure 5.**
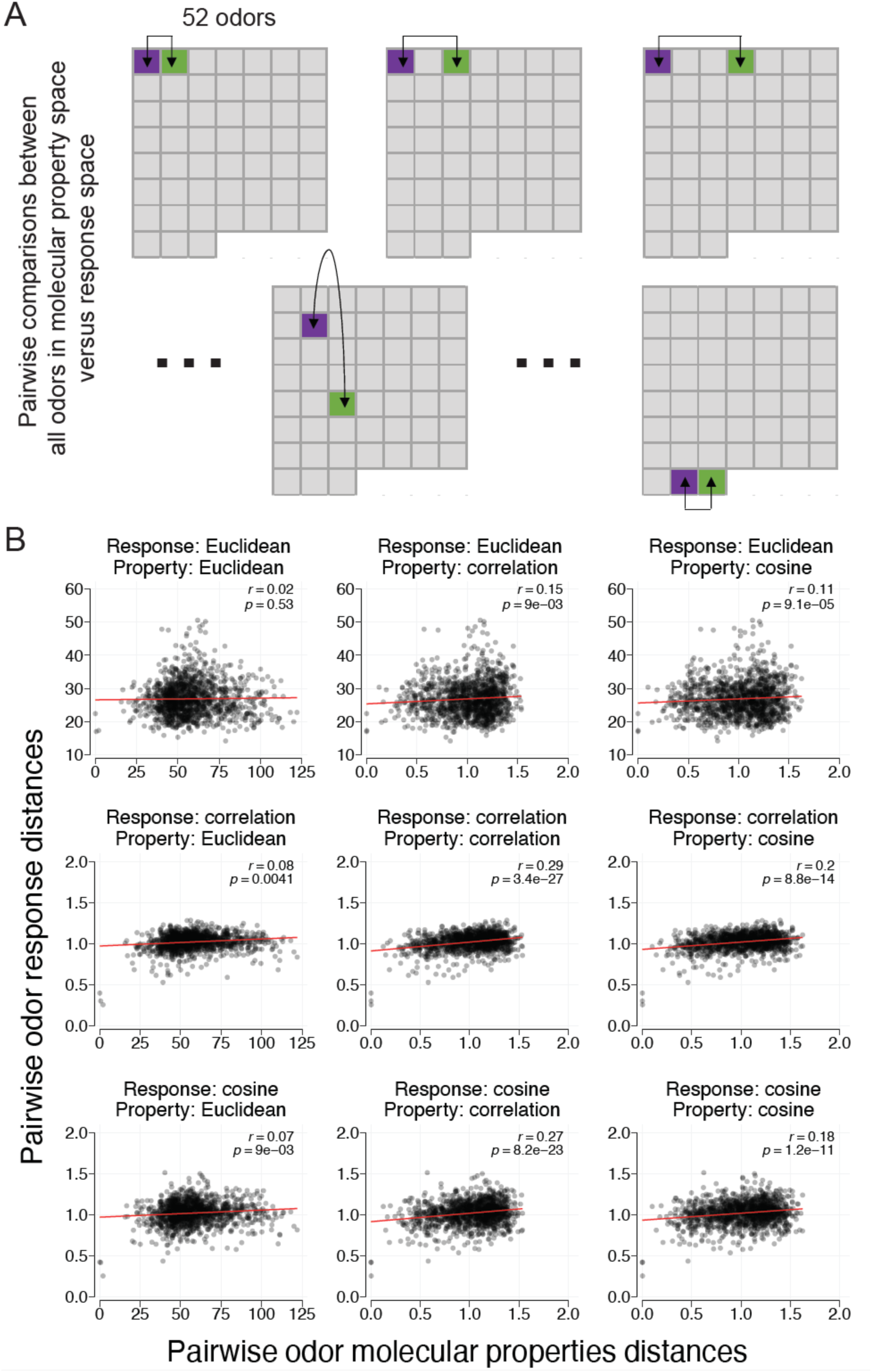
Details of the pairwise comparisons between odorants in response and molecular property space. **A**, Schematic of how pairwise distance comparisons between odorants were calculated for figures 3A and 3C. **B**, Correspondence between odorants in response and molecular property spaces. Three (Euclidean, correlation, and cosine) different distance metrics were used. Pearson correlation coefficients and *p*-values are reported for each combination of distance metric.

**Supplementary figure 6.**
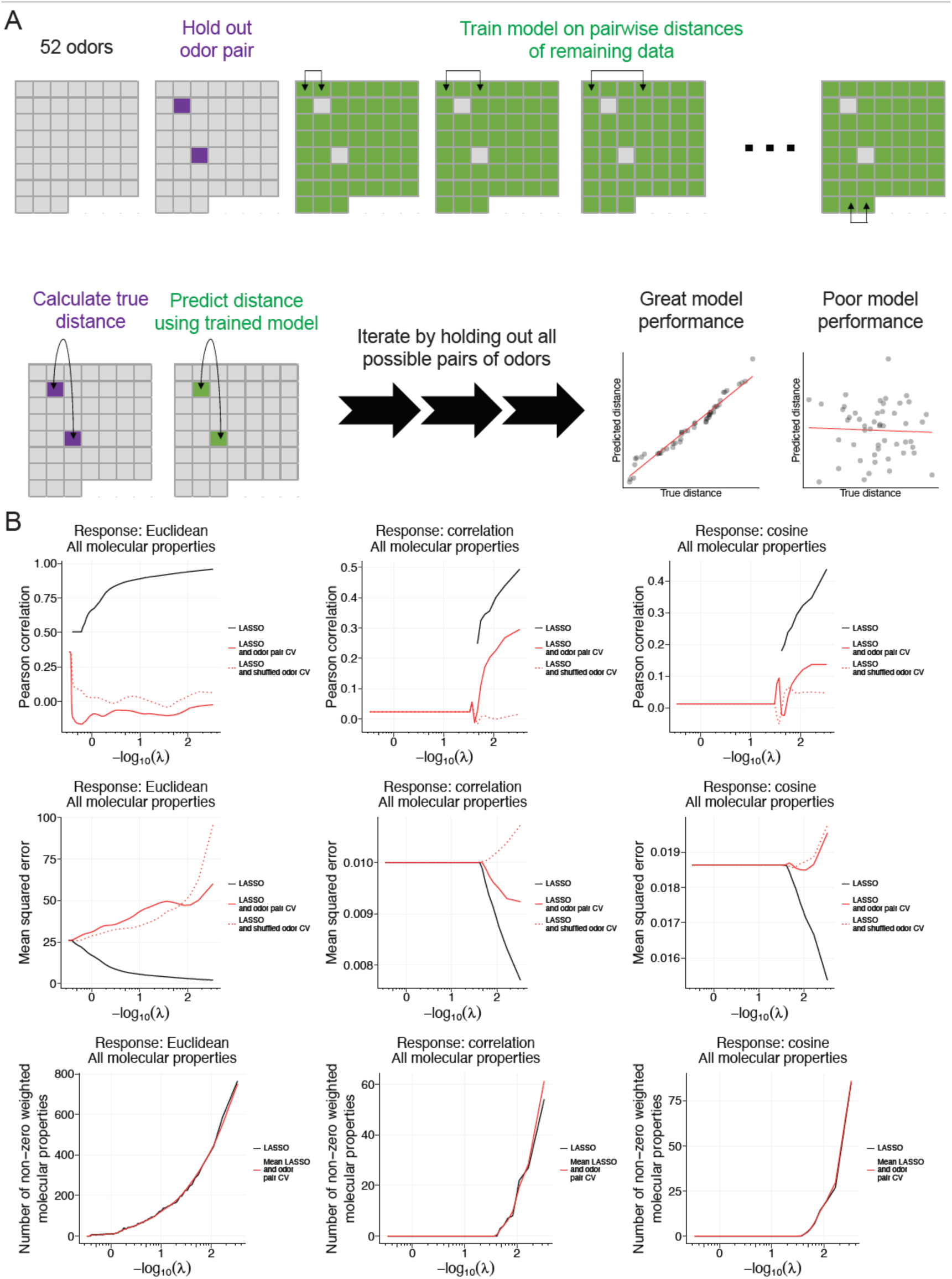
Details of the odor pair cross-validation for regression models. **A**, Schematic of how odor pair cross-validation was performed. Pairs of odorants were iteratively held-out from training. Distances between held-out odorants were iteratively predicted and regressed against true distances to report a Pearson correlation and mean squared error. **B**, Results of LASSO regression using various metrics to quantify distances between odorants in response space. Molecular property distances were quantified by Euclidean distances. Reported are Pearson correlation coefficients, mean squared error, and the number of non-zero weighted molecular properties as a function of varying the loss function (λ).

**Supplementary figure 7.**
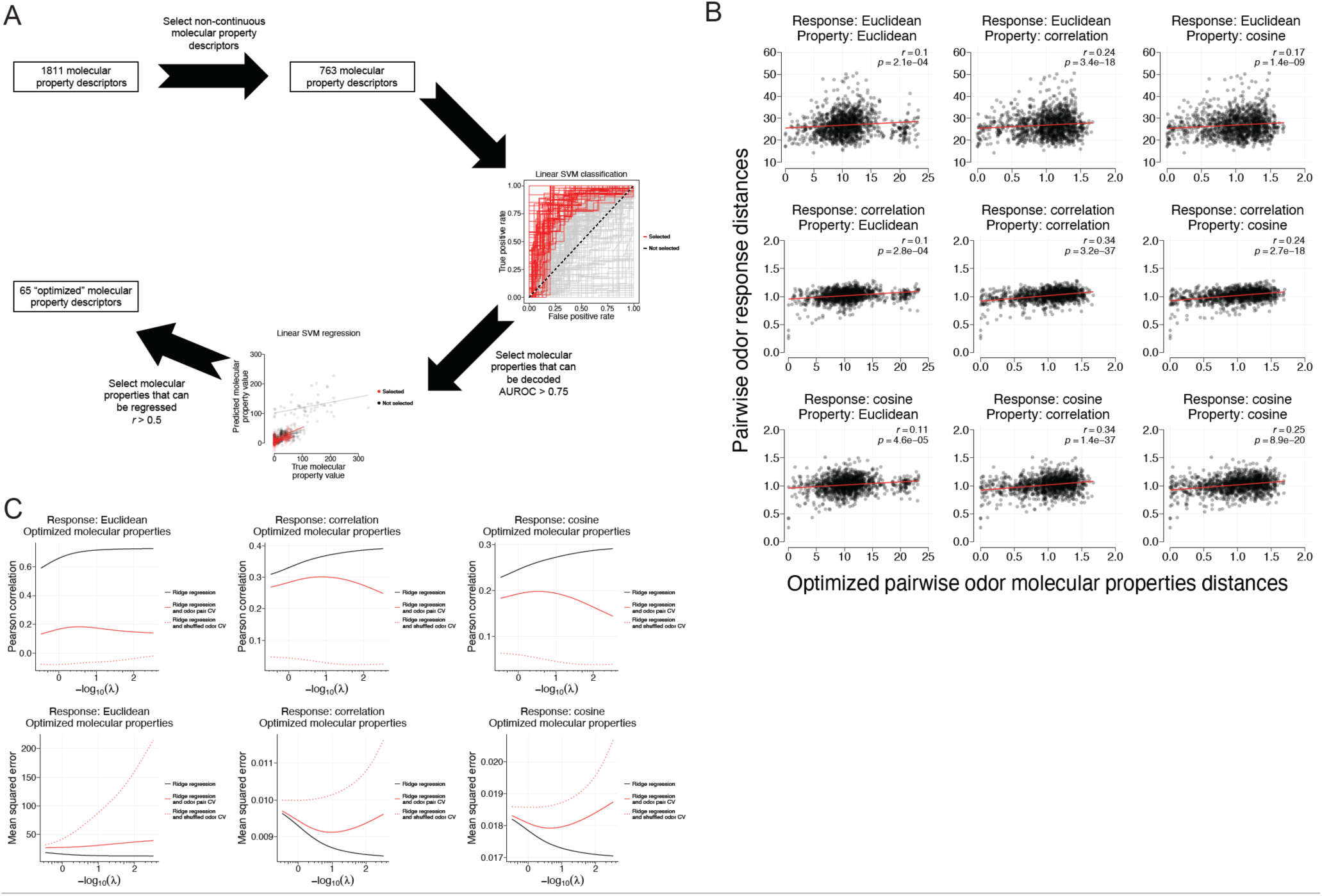
Performance of optimized molecular properties in predicting receptor response similarities. **A**, Schematic of how “optimized” molecular properties were selected. First, a set of 763 non-continuous descriptors were subset. Molecular properties that could be linearly decoded and regressed (with cross-validation) from the OR responses were then further selected using classifier AUROC thresholds of 0.75 and regressor *r* thresholds of 0.5. Ultimately 65 molecular properties passed these criteria and were considered “optimized”. **B**, Correspondence between odorants in response and “optimized” molecular property spaces. Euclidean, correlation, and cosine distance metrics are used to report Pearson correlation coefficients and *p*-values. **C**, Results of ridge regression using various metrics to quantify distances between odorants in response space. Generalizability was evaluated by odor pair cross-validation. Reported are Pearson correlation coefficients and mean squared error as a function of varying the loss function (λ).

**Supplementary figure 8.**
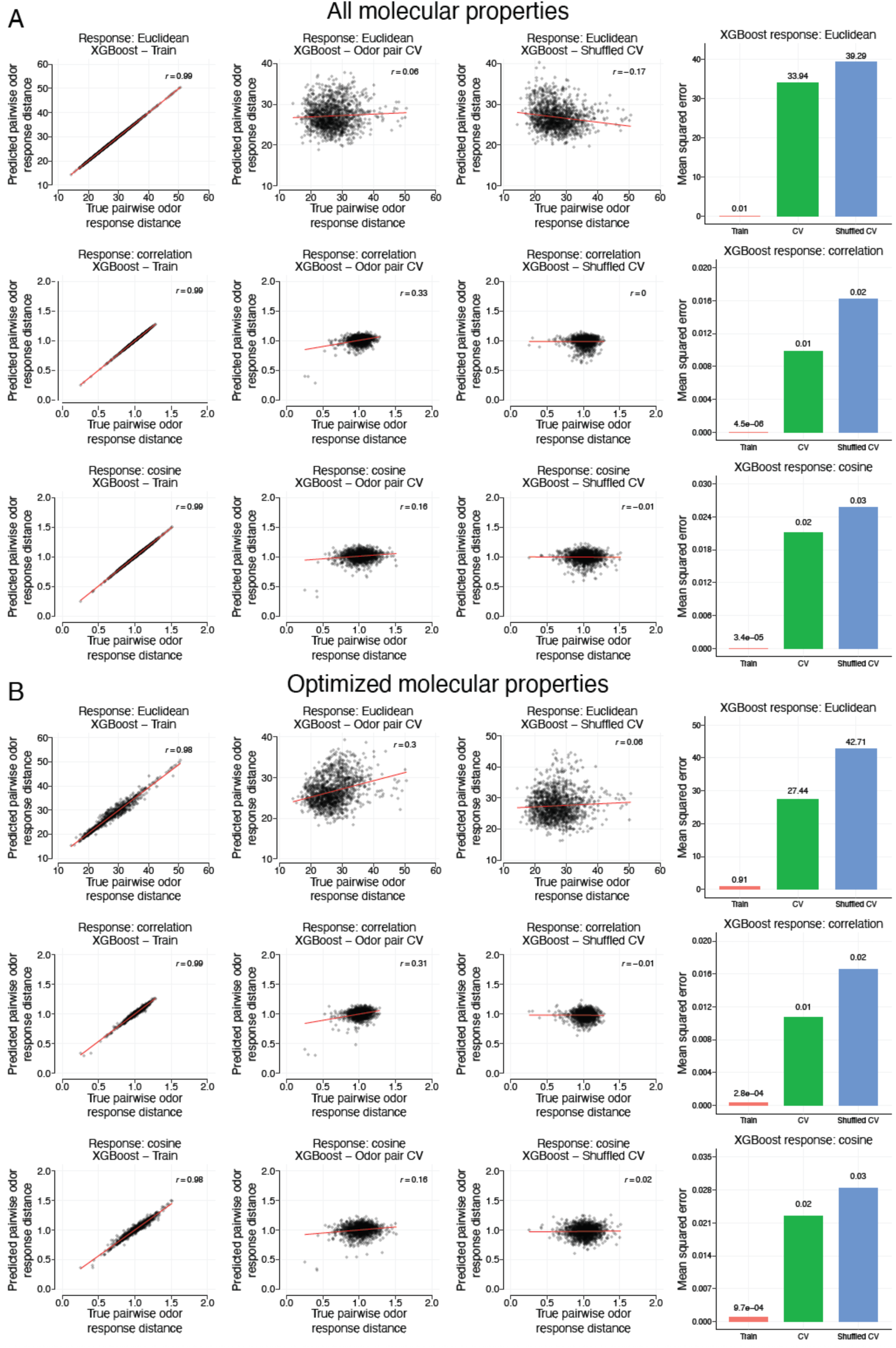
Results of using the XGBoost model framework. **A**, Using default XGBoost hyperparameters with 1811 molecular property descriptors, we asked how well does odor molecular property similarity predict receptor response similarity. Response similarities were calculated using Euclidean, correlation, and cosine distances. Odor pair cross-validation was performed to evaluate the generalizability of the models. **B**, Results of the XGBoost models when only the 65 set of “optimized” odor molecular properties were used as predictors and odor pair cross-validation is performed. The positive predictive abilities of these sparse 65 odor molecular properties, independent of the distance metric used) demonstrates odor similarity in response space can be approximated with parsimonious combinations of odor molecular properties.

**Supplementary figure 9.**
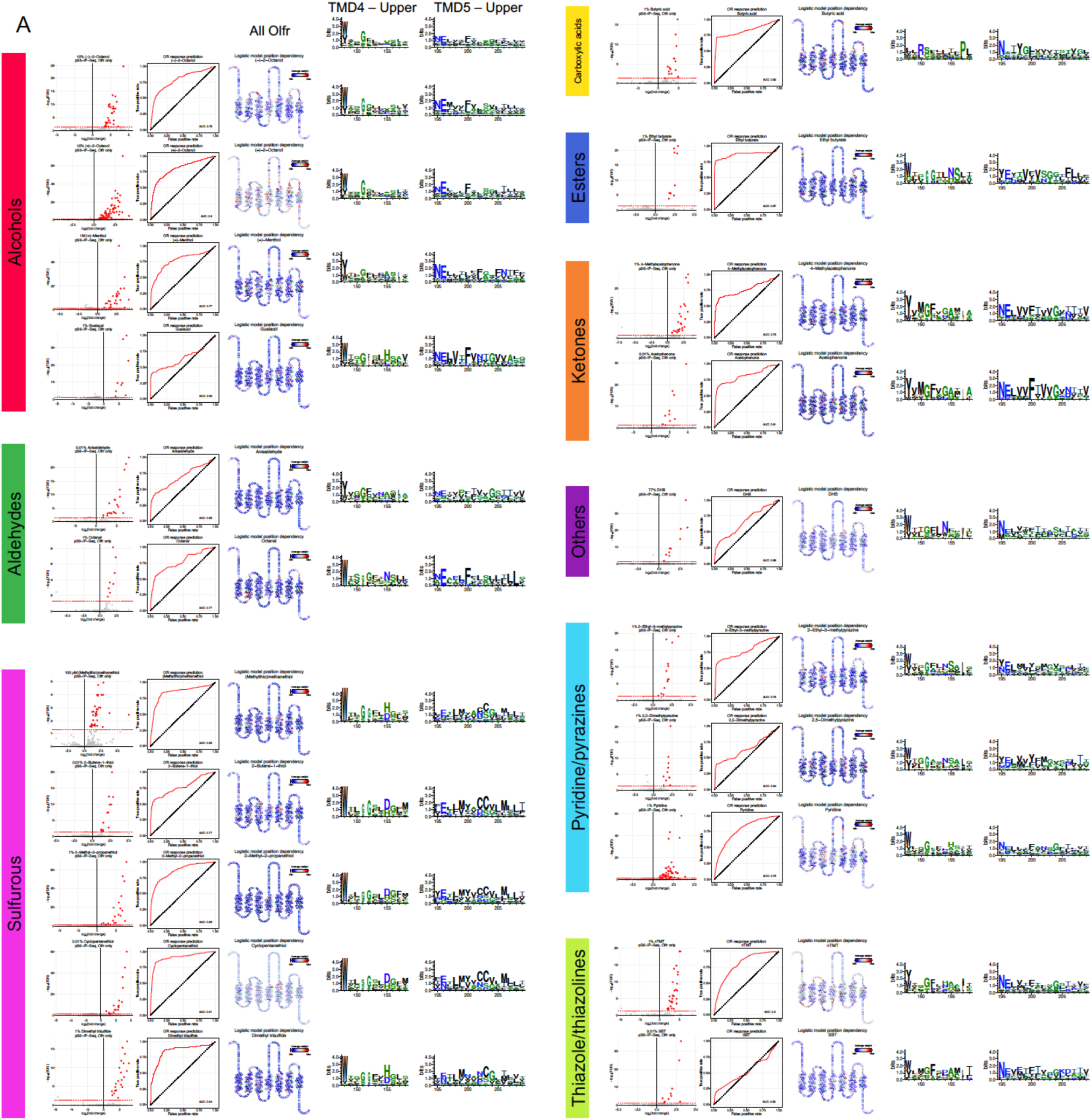
Details of logistic regression to uncover sequence-function relationships of ORs. **A,** Data for odorants solvable by regularized logistic regression (AUROC > 0.5) is shown. For each odor, shown is a volcano plot highlighting responsive ORs in red (log_2_FC > 0 and FDR < 0.05) and non-responsive ORs in gray. Aligned amino acid sequences were used as inputs to predict response likelihoods of a held-out 10% of ORs for 100 repetitions. The receiver operating characteristic (ROC) curve for held-out data is shown. Positions harboring amino acids assigned non-zero weights were averaged and visualized by snakeplot. Ballesteros-Weinstein x.50 numbers for each TMD are highlighted in green. The consistency of high weights assigned to residues localized to the upper halves of the fourth and fifth transmembrane domains motivated visualization of the amino acid distributions of responsive ORs by WebLogo. An enrichment of cysteine residues can be seen amongst ORs responsive to sulfurous odorants at positions 202 and 203 in TMD5. Similarly, an enrichment of methionine residues can be seen amongst ORs responsive to sulfurous odorants at positions 199 and 206.

**Supplementary figure 10.**
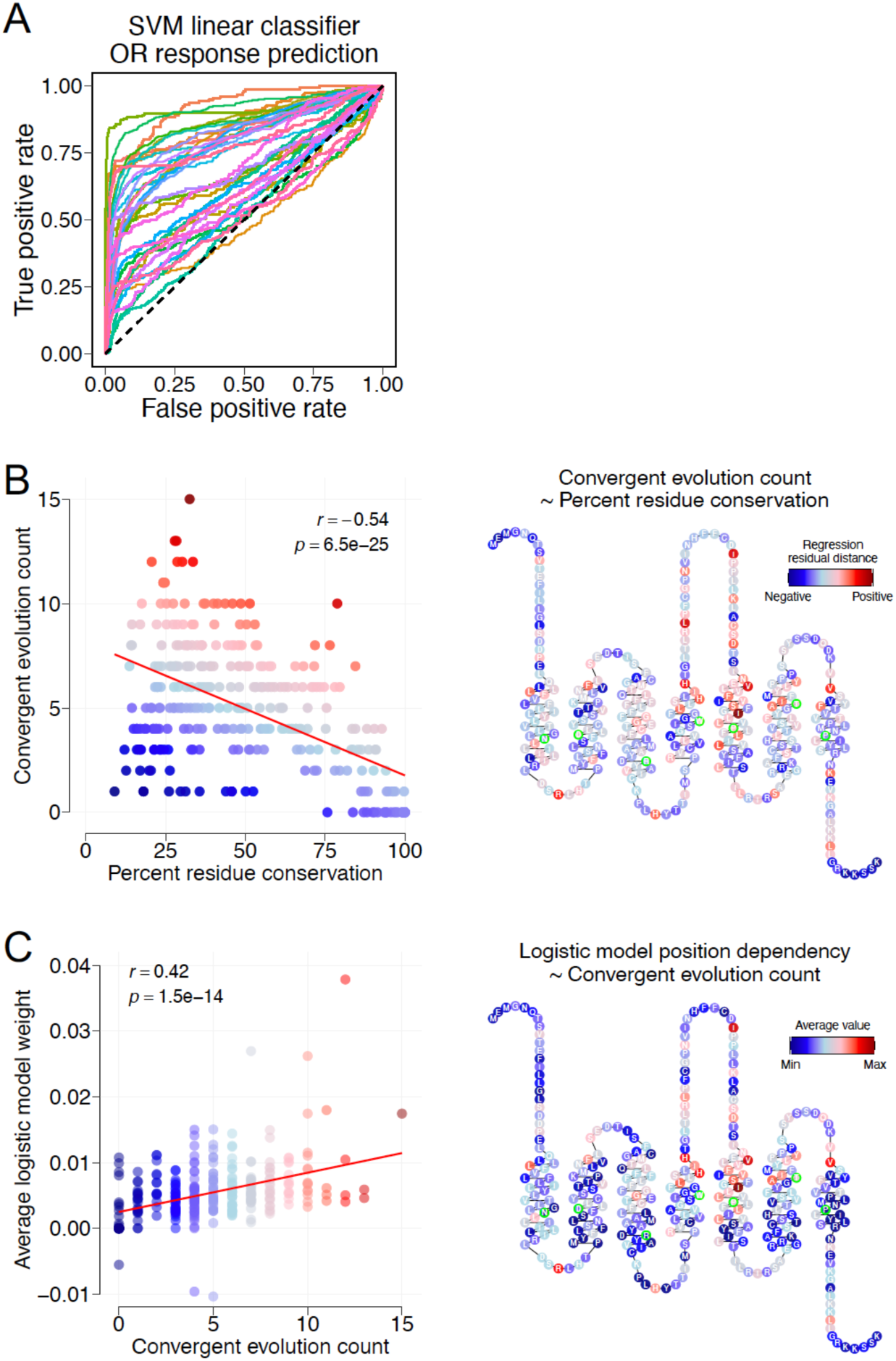
Details of SVM classifiers and comparing between approaches. **A,** Results of using a linear SVM classifier to predict OR response likelihoods across 100 repetitions of splitting ORs into 90% training and 10% testing sets. Models were trained and optimized using ten-fold cross-validation. Prediction likelihoods across 100 repetitions were compounded to generate single ROC curves for single odorants. AUROC values for each odorant are reported in supplementary table 4. **B,** Regressing convergent evolution counts by percent residue conservation reveals an anti-correlation (*r* = -0.54, *p* = 6.5E-25). Snakeplot is colored by the distance of each data point from the line of regression. Ballesteros-Weinstein x.50 numbers for each TMD are highlighted in green. **C,** Regressing logistic model dependency, evaluated by average positional weight, by convergent evolution count reveals consistency between approaches (*r* = 0.42, *p* = 1.5E-14).

## Methods

### Phosphorylated S6 ribosomal capture (pS6-IP)

Mice used for pS6-IP were ∼3 weeks old, mixed sex, and littermates. Mice were killed by CO_2_ asphyxiation and cervical dislocation. Olfactory tissue was rapidly dissected in Buffer B (2.5 mM HEPES KOH pH 7.4, 0.63% glucose, 100 μg/mL cycloheximide, 5 mM sodium fluoride, 1 mM sodium orthovanadate, 1 mM sodium pyrophosphate, 1 mM β-glycerophosphate, in Hank’s balanced salt solution). Tissue pieces were then minced in 1.35 mL Buffer C (150 mM KCl, 5 mM MgCl_2_, 10 mM HEPES KOH pH 7.4, 0.100 μM Calyculin A, 2 mM DTT, 100 U/mL RNAsin, 100 μg/mL cycloheximide, protease inhibitor cocktail, 5 mM sodium fluoride, 1 mM sodium orthovanadate, 1 mM sodium pyrophosphate, 1 mM β-glycerophosphate)and subsequently transferred to homogenization tubes for steady homogenization at 250 rpm three times and at 750 rpm nine times at 4 °C. Samples were then transferred to a 1.5 mL LoBind tube (Eppendorf 022431021) and clarified at 2000xg for 10 min at 4 °C. The low-speed supernatant was transferred to a new tube on ice, and 90 μL of NP40 (Sigma 11332473001) and 90 μL of 1,2-diheptanoyl-sn-glycero-3-phosphocholine (DHPC, Avanti Polar Lipids 850306P, 100 mg/0.69 mL) were added to this solution. This solution was mixed and then clarified at a max speed (17,000xg) for 10 min at 4 °C. The resulting high-speed supernatant was transferred to a new tube where 20 μL was saved and transferred to a tube containing 350 μL buffer RLT. To the remainder of the sample, 1.3 μL of 100 μg/mL cycloheximide, 27 μL of phosphatase inhibitor cocktail (250 mM sodium fluoride, 50 mM sodium orthovanadate, 50 mM sodium pyrophosphate, 50 mM β-glycerophosphate) and 6 μL of anti-pS6 antibody (Cell Signaling D68F8) were added. The sample was gently rotated for 90 min at 4 °C. To prepare beads, 100 μL of beads (Invitrogen 10002D) was washed three times with 900 μL of buffer A (150 mM KCl, 5 mM MgCl_2_, 10 mM HEPES KOH pH 7.4, 10% NP40, 10% BSA), and once with 500 μL of buffer C. Sample homogenate was added to the beads and incubated with gentle rotation for 60 min at 4 °C. Following incubation, beads were washed with four times with 700 μL of buffer D (350 mM KCl, 5 mM MgCl_2_, 10 mM HEPES KOH pH 7.4, 10% NP40, 2 mM DTT, 100 U/mL RNAsin, 100 μg/mL cycloheximide, 5 mM sodium fluoride, 1 mM sodium orthovanadate, 1 mM sodium pyrophosphate, 1 mM β-glycerophosphate). During the final wash, beads were moved to room temperature, wash buffer was removed, and 350 mL of buffer RLT was added. Beads were incubated in buffer RLT for 5 min at room temperature. Buffer RLT containing immunoprecipitated RNA was then eluted and stored at −80 °C until clean up using a kit (Qiagen 74004). cDNA was generated using 11 rounds of amplification with 10 ng RNA input. DNA libraries were prepared using a half-sized Nexterra XT DNA Library Preparation Kit (Illumina 15032354) protocol as per the manufacturer’s guidelines. Libraries were sequenced on either HiSeq 2000/2500 (50 base pair single read mode) or NextSeq 500 (75 base pair single read mode) with 6–12 pooled indexed libraries per lane.

### RNA-Seq alignment, quantification, and differential expression analysis

Reads were aligned against a modified GRCm38.p6 (M25) reference, in which we deleted ENSMUSG00000116179 (Olfr290), using STAR^39^ with --outFilterMultimapNmax 10. Reads mapping to Olfr290 were inferred from ENSMUSG00000070459, with the rationale that this gene model included ENSMUSG00000116179 plus untranslated regions. Gene-level read quantification was done using RSEM^40^. Differential expression analysis was performed against all genes using EdgeR^41^. Gene nomenclature was retrieved from BioMart^42^. Intact *Olfr* genes with identifiable sequences were filtered, and p-values were then re-corrected by FDR. Only ORs exhibiting odor response to at least one of the tested odorants (log_2_FC > 0 and FDR < 0.05) were considered. A total of 555 ORs responded across the 72 different odorants at various concentrations. A total of 375 ORs were responsive to unique odorants at the lowest tested concentrations. Raw and processed RNA-Seq datasets generated as part of this study are available from NCBI GEO at accession GSE185415.

### Source of odorants

The following odors/concentrations were used for comparing molecular properties to receptor responses: 1% 2-methyl-2-pentenal (Sigma 294667), 1% *trans*-cinnamaldehyde (Sigma C80687), 1% 2-heptanone (Sigma 537683^15^), 1% linalool (Sigma L2602), 1% ethyl butyrate (Sigma W242713), 1% guaiacol (Sigma G10903), 1% diacetyl (Sigma W237027), 1% 2-ethyl-3-methylpyrazine (Sigma W315508), 1% 2,5-dimethylpyrazine (Sigma 175420^15^), 1% benzaldehyde (Sigma W212717), 1% (+)-limonene (Sigma 183164), 1% β-damascone (Sigma W324300), 1% α-pinene (Sigma W290267), 1% 2- methyl-2-thiazoline (Sigma M83406), 1% citronellol (Sigma W230915), 1% dimethyl trisulfide (Sigma W327506), 1% *p*-Cresol (Sigma C85751), 0.01% citral (Sigma W230316), 1 M (+)-menthol (Sigma 224464), 1 M (-)-menthol (Sigma M2780), 0.01% anisaldehyde (Sigma A88107), 1% 4-methylacetophenone (Sigma W267708), 1% methyl salicylate (Sigma W274502), 1% (+)-carvone (Sigma 22070), 1% (-)-carvone (Sigma 22060), 1% β-ionone (Sigma W259525), 1% isopropyl tiglate (Sigma W322903), 1% hexyl tiglate (Sigma W500909), 1% pyridine (Sigma 270970), 1% butyric acid (Sigma W222119), 0.01% cyclopentanethiol (Sigma W326208), 0.01% 2-butene-1-thiol (1717 CheMall Corp OR116574), 100 mM cyclopentadecanone (Sigma C111201), 1% 2-methyl-2-propanethiol (Sigma 109207), 0.01% acetophenone (Sigma W200910), 0.1% isovaleric acid (Sigma 129542), 1% isoamyl acetate (Sigma 306967), 1% ethyl tiglate (Sigma W246000), 1% heptanoic acid (Sigma W334812), 10% (+)-2-octanol (Sigma O4504), 10% (-)-2-octanol (Sigma 147990), 1% 2-hexanone (Sigma 103004), 1% 2-phenylethanol (Sigma 77861), 1% 3-methyl-1-butanethiol (Sigma W385808), 1% octanal (Sigma O5608), 1% heptanal (Sigma W254002), 1% 2,4,5-trimethylthiazole (nTMT, Sigma 219185), 100% (E)-*β*-Farnesene (Bedoukian P3500-90^14^), 100 μM (methylthio)methanethiol (MTMT, synthesized^15^), 0.01% 2-*sec*-butyl-4,5-dihydrothiazole (SBT, synthesized^15^), 77% 3,4-dehydro-*exo*-brevicomin (DHB, synthesized^15^), and 0.01% 2,4,5-trimethyl-4,5-dihydrothiazole (TMT, synthesized^14^).

For logistic regression and identifying residues with predictive power towards ligand selectivity, odorants tested at the lowest concentration with at least 8 activated ORs (log_2_FC > 0 and FDR < 0.05) were used to promote class stability. Thus, following odors were removed from consideration using logistic regression compared to above: 100 mM cyclopentadecanone, 1% 2-heptanone, 1% 2-hexanone, 1% 3-methyl-1-butanethiol, 1% *α*-pinene, 1% benzaldehyde, 1% *β*-ionone, 1% ethyl tiglate, 1% heptanoic acid, 1% hexyl tiglate, 1% isopropyl tiglate, 1% linalool, 1% methyl salicylate, 1% (+)- limonene, 1% *trans*-cinnamaldehyde, and 0.1% isovaleric acid. The following odors were considered at a modified concentration from above: 0.1% TMT, 0.1% acetophenone, and 10 mM MTMT.

Odorants were excluded from all analysis if no ORs were identified as responsive at the tested concentrations: 1% β-Caryophyllene (Sigma W225207^15^), 1% dimethyl sulfide (Sigma 274380), 1% geraniol (Sigma W250716), 1% indole (Sigma W259378), 1% (-)-dihydrocarveol (Sigma 37278), 1% (+)-dihydrocarveol (Sigma 37277), 1% propionic acid (Sigma 109797), 3mM androstenone (Sigma 284998), or if the number of ORs identified as responsive were more than five standard deviations away from the mean: 1% 2’-hydroxyacetophenone (Sigma H18607).

### Chemical space estimation

To estimate chemical space, we first identified 4680 small molecules commonly found in foods and fragrances from http://www.thegoodscentscompany.com/^16^. Three dimensional structures for these molecules and the 52 in the test odor set were then downloaded from PubChem, and 5666 molecular properties were calculated using AlvaDesc (v2.0.10). From the 5666 calculated molecular properties, 3855 were discarded because they were either not calculated for all molecules or exhibited zero variance across all molecules, leaving behind 1811 molecular descriptors. Chemical space was estimated by PCA dimensionality reduction on all molecules in R.

### Receptor alignment and space estimation

Mouse ORs were aligned to one another using the MAFFT E-INS-I method with manual refinements^43^. The resulting alignment file was subjected to ModelTest-NG to identify ideal amino acid substitution models^44^. Phylogenetic trees were generated with RAxML-NG using the JTT+I+G4 amino acid substitution model with 100 bootstraps^45^. Receptor pairwise similarity matrices for multidimensional scaling were generated from an alignment in which positions with amino acids in at least 60% of the receptors were considered. Receptor pairwise similarity was calculated by summing amino acid differences at each position by Grantham’s amino acid distances^17^. Multidimensional scaling was done in R.

### Generating response spectra

To generate odor response spectra, we first began with the log_2_FC values of each odor-responsive OR. Each OR, *r*, was then centered and scaled (z-scored) by mean subtraction and standard deviation division across the odorants, *o*, in the test panel. The resulting matrix is denoted as 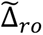. To generate property strength vectors, each molecular property, *p*, was z-scored across the odorants in the test panel. The resulting matrix is denoted as 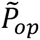. To calculate property responses and thereby property response spectra (Pearson correlation coefficients), we used the following formula:

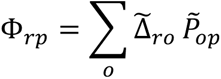

where Φ*_rp_* refers to Pearson correlation coefficients between individual receptors, *r*, and molecular properties, *p*.

To evaluate the significance of the correlation between a property and the response pattern of an OR, we used an FDR cutoff of 0.05. *P*-values were obtained by first calculating the t-statistic using 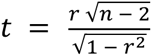, where *r* is the correlation coefficient and *n* is the number of data points. The two-tailed *P*-value was then calculated as twice the probability a *t*-distributed variable exceeds *t* using the python scipy.stats.t.sf function. *P*-values were adjusted by FDR correction in R.

### Odor distance calculation in property space and response space

We calculated the distances between odorants in property and response space by calculating Euclidean distances, Pearson correlation coefficients, and cosine similarities between all possible unique pairs of odorants. Molecular properties and receptor response data were normalized to their respective zero mean and unit standard deviation. Correlation distances were reported as 1 – *r*, and cosine distances were reported as 1 – *cos(θ)*.

### Regression models with odor pair cross-validation

Linear models (LASSO and ridge regression) were implemented with the glmnet package (v4.1) in R^46^. XGBoost was implemented with the xgboost module (v1.4.2) in python. Distances (Euclidean, cosine, and correlation) between each unique pair of odorants were calculated in normalized receptor response space. Then, Euclidean distances between each unique pair of odorants were calculated for each feature in normalized feature space. Regularization was then applied as either the L1 (LASSO) or L2 (ridge regression) norm. The λ loss function, which controls the number and relative contribution of selected features, was sequentially varied from zero to three by length 1000. Pearson correlation values were reported for the varied λ hyperparameters of the models by comparing model predicted response distances to true response distances. Shuffled controls consisted of using 52 fictitious odorants whose individual feature vectors were generated by resampling with replacement across the 52 test odorants.

Default XGBoost model hyperparameters were used as follows: base_score=0.5, booster=’gbtree’, colsample_bylevel=1, colsample_bynode=1, colsample_bytree=1, gamma=0, importance_type=’gain’, interaction_constraints=’’, learning_rate=0.300000012, max_delta_step=0, max_depth=6, min_child_weight=1, missing=nan, monotone_constraints=’()’, n_estimators=100, num_parallel_tree=1, random_state=42, reg_alpha=0, reg_lambda=1, scale_pos_weight=1, subsample=1, tree_method=’exact’, validate_parameters=1, and verbosity=None.

Odor pair cross-validation was performed by iteratively holding out each unique pair of odorants from the normalized 52 odor dataset (test data). Features with zero variance from the remaining 50 odor set were dropped (train data). Distances were calculated between the test data for each remaining feature (xtest) and response pattern (ytest). Train data were normalized by z-scoring independent of test data. Distances were then calculated between pairwise combinations of the 50 train odorants for each feature (xtrain) and response pattern (ytrain). Feature distances between the held-out odor pair (xtest) were then used to predict response pattern distances (ypred). Pearson correlation and mean squared error values were reported from comparing model predicted response distances (ypred) to true distances (ytest).

### Optimized molecular property selection

To select a subset of molecular properties that were well represented in the data, we utilized Support Vector Machine (SVM) classifiers and regressors with linear kernels in the python sklearn.svm module (v0.24.2). Beginning with the 1811 molecular properties, we first considered those that were non-continuous (at least one zero entry, ex. molecular weight is a continuous molecular property). Non-zero values were set to one and zero values were kept. Classifiers were trained and cross-validated across the 52 odorant molecules using the normalized 375 deorphanized receptor responses as predictors in a leave-one-out scheme. Data normalization was first performed including the test data. After removing test data, training data were normalized independently to prevent contamination. Classifier area under receiver operating characteristic (AUROC) thresholds of 0.75 were applied. Molecular properties passing this threshold were next subjected to regression with non-zero entries restored. A Pearson correlation cutoff of 0.5 was applied to finally select the 65 “optimized” molecular properties.

### Protein sequence analysis of ORs by logistic regression and SVM classifiers

Regularized logistic regression was used to build models linking OR-protein sequence properties to OR-odor responses with the glmnet package (v4.1) in R^46^. ORs were classified as responders if they exhibited log_2_FC > 0 and FDR < 0.05 following pS6-IP-Seq and differential expression analysis. For odorants tested at multiple concentrations, the lowest concentration that activated at least 8 ORs was used to promote class stability. Predictors were generated from converting the FASTA alignment file into categorical variables reflecting the presence/absence of specific amino acids at each position.

To evaluate model performance, fitted odorants were randomly split into 90% training and 10% testing receptor sets. Each test set contained at least one responding receptor. Predictors with zero variance in the training set were dropped. The grid-search optimized α hyperparameter (setting the ratio of the L1 and L2 norms) was set by ten-fold cross-validation with ten-fold cross-validation to set the λ (loss function) value. λ values one standard error of mean greater than optimal were selected to encourage statistically identical but sparser solutions. Model weighted predictors were then used to determine the response likelihood of the test receptors. This procedure was repeated 100 times. Non-zero weights were averaged across repetitions and odorants to report positions with residues contributing predictive power towards odor selectivity. WebLogo visualizations were prepared at http://weblogo.threeplusone.com/47.

SVM classifier response probabilities were calculated using the same inputs as logistic regression using 100 repetitions of 90% training (with ten-fold cross-validation for hyperparameter tuning) and 10% testing. Each repetition’s response likelihoods and true outcomes were aggregated to generate a single ROC curve for a single odor, which were then combined to generate an aggregate ROC curve.

### Protein sequence analysis of ORs by comparison to convergently evolved ORs

As an alternative strategy, we also performed a statistical evaluation of amino acid properties of ORs sharing responsiveness to an odor against convergently evolved receptors. First, responsive ORs (log_2_FC > 0 and FDR < 0.05 from differential expression) were subset, and pairwise Grantham distances were calculated at each position to generate Grantham distance distributions within the responsive OR alignment. Pairwise comparisons between gaps were considered to have zero distance while pairwise comparisons between gaps and amino acids were considered to have the average Grantham distance across all pairwise comparisons between all ORs at that position. Null distributions were generated similarly from convergently evolved odor-unresponsive ORs. To identify convergently evolved odor-unresponsive sets of ORs, odor-specific receptors with log_2_FC < 0 or FDR > 0.25 were first subset. Then, for each unique pairwise comparison between the odor-responsive ORs, full protein sequence Grantham distances were calculated. For each receptor in each pairwise comparison, the closest receptor was selected from the odor-unresponsive subset with the most similar absolute full protein sequence Grantham distance to the pairwise comparison. This meant, for each odor with some number of responsive receptors, there was twice as many receptors identified as convergently evolved and odor-unresponsive. Distributions were compared using the Kolmogorov-Smirnov statistical test. FDR correction was applied across all calculated *P*-values with a cutoff of 0.05. The number of times responding receptors displayed statistically significant deviations in the distribution of Grantham distances from the null set, at each position, was counted and summed across all odorants.

### Residue conservation calculation

Using the 313 length alignment file, in which each position was occupied by an amino acid in at least 60% of the responsive ORs (387 ORs that were responsive to the lowest concentration of tested odorants yielding response to at least 8 ORs each), we first identified the most common amino acid at each position. We term this the reduced consensus OR sequence. The percent presence of the most commonly occurring amino acid at each position was then reported as conservation percentage for said position.

### Homology models

To build an OR homology model, we adapted previously published methods^48, 49^. The reduced consensus OR sequence was manually re-aligned to pre-aligned sequences of the bovine rhodopsin (PDB ID 1U19), the human chemokine receptors CXCR4 (3ODU) and CXCR1 (2LNL), and human adenosine A2A receptor (2YDV) using Jalview. Experimental GPCR structures of these receptors were then used as templates to build the homology model of the reduced consensus sequence with Modeller. Visualization and analysis of the homology model was done using VMD and Chimera.

### Heterologous luciferase assay

Hana3A cells; which stably express G_olf_, RTP1, RTP2, and REEP1; were grown in minimum essential medium eagle (MEM; Corning 10-010-CV) containing 10% Fetal Bovine Serum (FBS; vol/vol; Gibco 16000-044), penicillin-streptomycin (Sigma-Aldrich P4333), and amphotericin B (Gibco 15290018). Cells were cultured and incubated at 37°C, 5% CO_2_, and saturated humidity for use with the Dual-Glo Luciferase Assay (Promega E2980)^29, 50^. Cells were plated at 20-25% confluence on poly-D-lysine-coated 96-well plates (Corning 3843) overnight. After overnight incubation, cells were transfected with 6 mL of MEM containing 10% FBS, 0.5 µg SV40-RL (Promega E2980), 1 µg CRE-Luc (Promega E2980), 0.5 µg mouse RTP1s, 0.25 µg M3 muscarinic receptor^51^, 0.5 µg of Rho-tagged receptor plasmid DNA, and 20 µg Lipofectamine 2000 (Invitrogen 11668019) per plate. Transfection medium was divided equally among the wells so that each OR-odorant combination could be conducted in triplicates. The following day, cells were incubated with 25µL of odorant solution diluted in CD-293 (Gibco 11913-019) containing 30 µM CuCl_2_ (Sigma-Aldrich C-6641) and 2 mM glutamine (Gibco 25030-081) for 3.5 hours. cAMP-driven firefly Luciferase luminescence (Luc) was used to assess OR activation, and SV40-driven *Renilla* Luciferase luminescence (Ren) was used to control for variation in cell viability within wells. Cell luminescence was read by a POLARstar OPTIMA (BMG Labtech) luminometer, and normalized response values were calculated using the formula (Luc-400)/(Ren-400). ORs were considered responsive *in vitro* if ANOVA *p*-value was < 0.05 and ANOVA with post-hoc Dunnet’s test correction *p*-adjusted was < 0.05 for at least 2 of the tested odor concentrations using the R package DescTools (v0.99.42). Log-logistic 4-parameter dose response curves were fit to the data using the R package drc (v3.0-1). *In vitro* responses were compared to *in vivo* responses by subtracting mean ligand-independent activity (luciferase values of ORs with no odor stimulation) from each of the ligand stimulated data points and summing. Scaled summed (+)-enantiomer responses were divided by scaled summed (-)-enantiomer responses and log_2_ transformed for comparison to log_2_FC (+)/(-) *in vivo* enrichments.

## Data and code availability

Data and code are available upon reasonable request.

## Contributions

AV, MHN, XSH, and HM did *in vivo* experiments. CJ and CAdM did *in vitro* experiments. AV analyzed data. CAdM did OR homology modeling. JP advised data modeling analysis. AV drafted the paper. All authors reviewed and edited the paper. HM supervised the work.

## Acknowledgements

We thank Michael Schmuker for critical reading and comments, Mengjue Jessica Ni for expert technical assistance, Alex Koulakov for helpful discussions on data analysis, Priya Meesa for manuscript edits, and members of the Matsunami lab for helpful discussions. AV thanks Jack J. Zhan for advising how to analyze data in python during initial stages.

